# On the Origin and Evolutionary Consequences of Gene Body DNA Methylation

**DOI:** 10.1101/045542

**Authors:** Adam J. Bewick, Lexiang Ji, Chad E. Niederhuth, Eva-Maria Willing, Brigitte T. Hofmeister, Xiuling Shi, Li Wang, Zefu Lu, Nicholas A. Rohr, Benjamin Hartwig, Christiane Kiefer, Roger B. Deal, Jeremy Schmutz, Jane Grimwood, Hume Stroud, Steven E. Jacobsen, Korbinian Schneeberger, Xiaoyu Zhang, Robert J. Schmitz

## Abstract

In plants, CG DNA methylation is prevalent in the transcribed regions of many constitutively expressed genes (“gene body methylation; gbM”), but the origin and function of gbM remain unknown. Here we report the discovery that *Eutrema salsugineum* has lost gbM from its genome, the first known instance for an angiosperm. Of all known DNA methyltransferases, only CHROMOMETHYLASE 3 (CMT3) is missing from *E. salsugineum*. Identification of an additional angiosperm, *Conringia planisiliqua*, which independently lost CMT3 and gbM supports that CMT3 is required for the establishment of gbM. Detailed analyses of gene expression, the histone variant H2A.Z and various histone modifications in *E. salsugineum* and in *Arabidopsis thaliana* epiRILs found no evidence in support of any role for gbM in regulating transcription or affecting the composition and modifications of chromatin over evolutionary time scales.

## Significance Statement

DNA methylation in plants is found at CG, CHG, and CHH sequence contexts. In plants, CG DNA methylation is enriched in the transcribed regions of many constitutively expressed genes (“gene body methylation; gbM”) and show correlations with several chromatin modifications. Contrary to other types of DNA methylation, the evolution and function of gbM is largely unknown. Here we show two independent concomitant losses of the DNA methyltransferase CMT3 and gbM without the predicted disruption of transcription and of modifications to chromatin. This result suggests that CMT3 is required for the establishment of gbM in actively transcribed genes, and that gbM is dispensable for normal transcription as well as the composition and modifications of plant chromatin.

In angiosperms, cytosine DNA methylation occurs in three sequence contexts: methylated CG (mCG) is catalyzed by METHYLTRANSFERASE I (MET1), mCHG (where H = A/C/T) by CHROMOMETHYLASE 3 (CMT3), and mCHH by DOMAINS REARRANGED METHYLTRANSFERASE 2 (DRM2) or CHROMOMETHYLASE 2 (CMT2) (1). MET1 performs a maintenance function and it is targeted by VIM1, which binds preexisting hemi-methylated CG sites. In contrast, DRM2 is targeted by RNA-directed DNA methylation (RdDM) for the *de novo* establishment of mCHH. CMT3 forms a self-reinforcing loop with the H3K9me2 pathway to maintain mCHG; however, considering that transformation of CMT3 into the cmt3 background can rescue DNA methylation defects, it is reasonable to also consider CMT3 a *de novo* methyltransferase (2). Two main lines of evidence suggest that DNA methylation plays an important role in the transcriptional silencing of transposable elements (TEs): that TEs are usually methylated, and that the loss of DNA methylation (e.g. in methyltransferase mutants) is often accompanied by TE reactivation.

A large number of plant genes (e.g. ~13.5% of all *Arabidopsis thaliana* genes) also contain exclusively mCG in the transcribed region and a depletion of mCG from both the transcriptional start and stop sites (referred to as “gene body DNA methylation”, or “gbM”) (**Fig. 1A**) (3–5). A survey of plant methylome data showed that the emergence of gbM in the plant kingdom is specific to angiosperms (6), whereas non-flowering plants (such as mosses and green algae) have much more diverse genic methylation patterns (7, 8). Similar to mCG at TEs, the maintenance of gbM requires MET1. In contrast to DNA methylation at TEs, however, gbM does not appear to be associated with transcriptional repression. Rather, genes containing gbM are ubiquitously expressed at moderate to high levels compared to non-gbM genes (4, 5, 9), and within gbM genes there is a correlation between transcript abundance and methylation levels (10, 11).

**Fig. 1.**
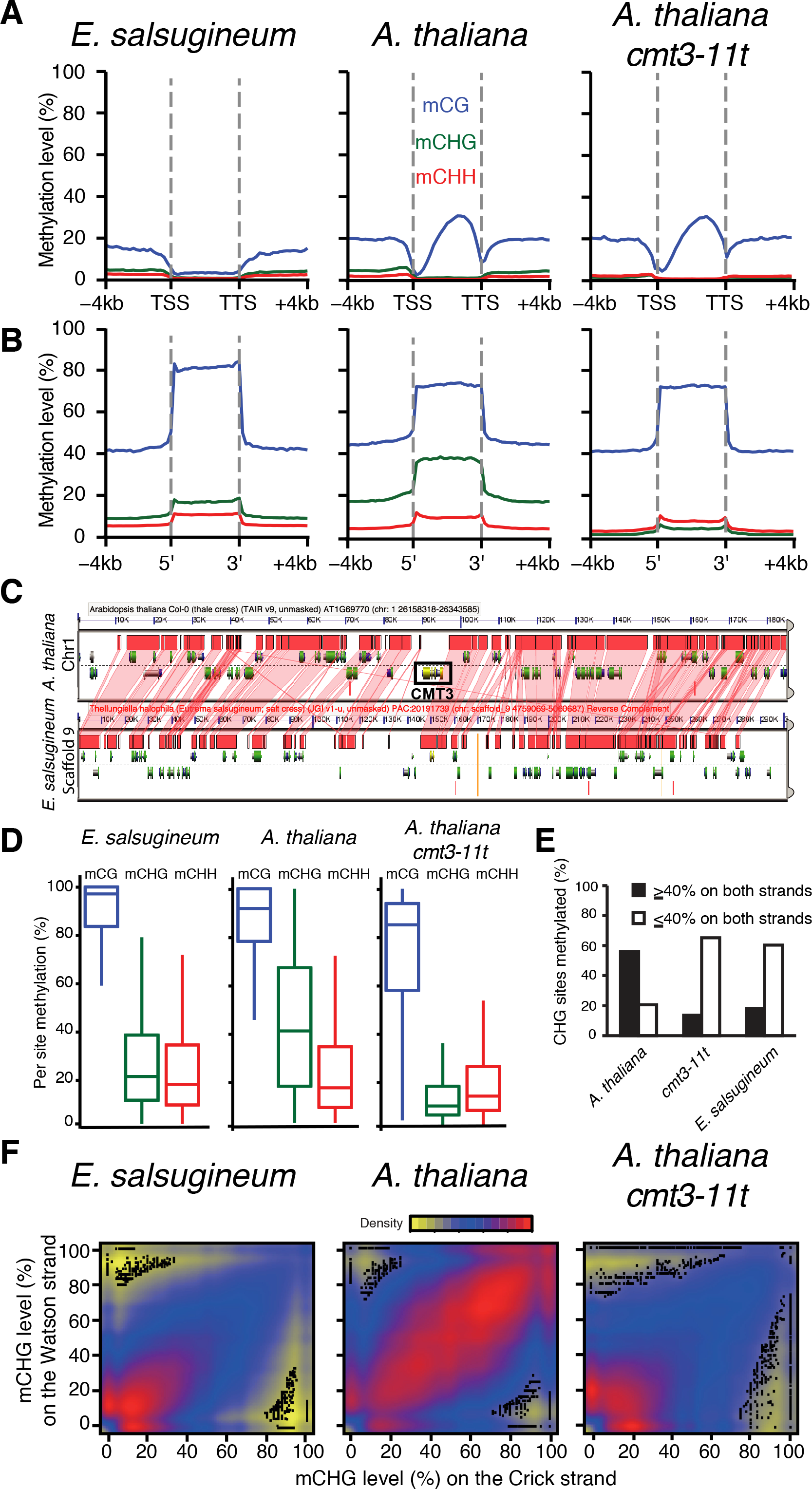
CMT3 and gbM is absent in *E. salsugineum.* Metagene plots of DNA methylation across **(A)** gene bodies and **(B)** repeats including 4 kb up-and down-stream. **(C)** A syntenic block of sequence is shared between *A. thaliana* and *E. salsugineum.* The black box in the *A. thaliana* block indicates the location of CMT3, which is absent in *E. salsugineum.* The red shaded areas indicate regions of shared synteny. **(D)** Boxplot representation of methylation levels of individual methylated cytosines within each sequence context. **(E)** A bar plot of methylation levels at symmetric CHG methylated sites in *A. thaliana, cmt3–11t* and *E. salsugineum.* **(F)** Density plot representation of CMT3-dependent versus CMT3-independent CHG methylation.

It has been proposed that gbM may be established by the *de novo* methylation activity of the RdDM pathway, and subsequently maintained by MET1 independently of RdDM. In this “*de novo*” scenario, occasional antisense transcripts could form double-stranded RNA by pairing with sense transcripts, which could trigger the production of small interfering RNAs (siRNAs) to target DRM2 for *de novo* methylation in gene bodies. Although mechanistically feasible, it is difficult to explain why gbM is absent from many non-angiosperm plants (such as the moss *Physcomitrella patens*) with functional RdDM and MET1 pathways (12).

Alternatively, we propose the establishment of gbM might involve the self-reinforcing loop between CMT3 and the histone H3 lysine 9 (H3K9) methyltransferase KRYPTONITE/SUVH4 (KYP) (13, 14) in addition to transcription, similar to a model proposed by Inagaki and Kakutani (15). CMT3 is recruited to chromatin by H3K9me2 for DNA methylation, which in turn recruits KYP for H3K9me2. Although mCHG and H3K9me2 are normally limited to heterochromatin, they accumulate ectopically in thousands of actively transcribed genes upon the loss of the H3K9 demethylase INCREASED IN BONSAI METHYLATION 1 (IBM1) (16). It therefore appears likely that mCHG and H3K9me2 also occur constantly (albeit transiently) in actively transcribed genes, but their accumulation is normally prevented by IBM1 (17). The transient presence of H3K9me2 in transcribed regions could trigger CMT3-dependent methylation of CHG and other contexts. The MET1 pathway would then maintain rare methylation of CG sites in an H3K9me2-independent manner. Consistent with this possibility, gbM-containing genes are preferred targets for hypermethylation in the *ibm1* mutant (16). Lastly, in support of a role for CMT3 in this model, CMT3 is only present in angiosperms, which coincides with the emergence of gbM in the plant kingdom (6, 17).

The difficulty in addressing the origin of gbM is two fold. First, gbM is unaffected by the loss of RdDM or CMT3 in the short term, indicating that the maintenance activity of MET1 is sufficient for the persistence of gbM over extended periods of time. Second, once gbM is lost in the *met1* mutant, it does not immediately return when MET1 is reintroduced by crossing, indicating that the establishment of gbM is a stochastic process that requires many generations (18).

Here we describe the results from a comparative epigenomics approach, where we sought to identify natural variation in plant methylomes (**Table S1**) that were associated with genetic changes in key genes in DNA methylation pathways. The methylomes of the vast majority of the plant species are similar to *A. thaliana*, with high levels of mCG/mCHG/mCHH colocalized to repetitive sequences, and gbM in moderately expressed genes (19). The only exception was *Eutrema salsugineum* (acc. Shandong), a member of the Brassicaceae family that shares a common ancestor with *A. thaliana* and *Brassica spp*. approximately 47 and 40 million years ago, respectively (20). Comparisons between the *E. salsugineum* methylome to those of other plants revealed two major differences. First, *E. salsugineum* has lost gbM (**Fig. 1A**). In contrast to other species where thousands of active genes contained gbM (e.g., 4,934 in *A. thaliana*), only 103 *E. salsugineum* genes contained gbM based on our identification criteria (**Dataset S1** - see methods). A closer inspection of these 103 loci revealed that the distribution of mCG in these loci was not representative of gbM genes in other angiosperms (**Fig. 2**). Importantly, mCG was present at high levels in repetitive sequences in *E. salsugineum*, indicating that the absence of gbM in its genome was not due to the loss of MET1 activity (**Fig. 1B**). Second, the mCHG level in *E. salsugineum* repetitive sequences was much lower compared to other plant species (**Fig. 1B**). To further validate these results, we performed MethylC-seq using an additional *E. salsugineum* accession (Yukon), and the results showed that it too has lost gbM (**Fig. 2** and Fig. S1). A detailed analysis of *E. salsugineum* genes identified homologs of all known DNA methyltransferases in *A. thaliana* (e.g. MET1, CMT2 and DRM2), with the exception of CMT3. In addition, a comparison of the genomic region between *E. salsugineum* syntenic to the *A. thaliana* CMT3 locus found no evidence for CMT3-related sequences at the syntenic location (**Fig. 1C**).

**Fig. 2.**
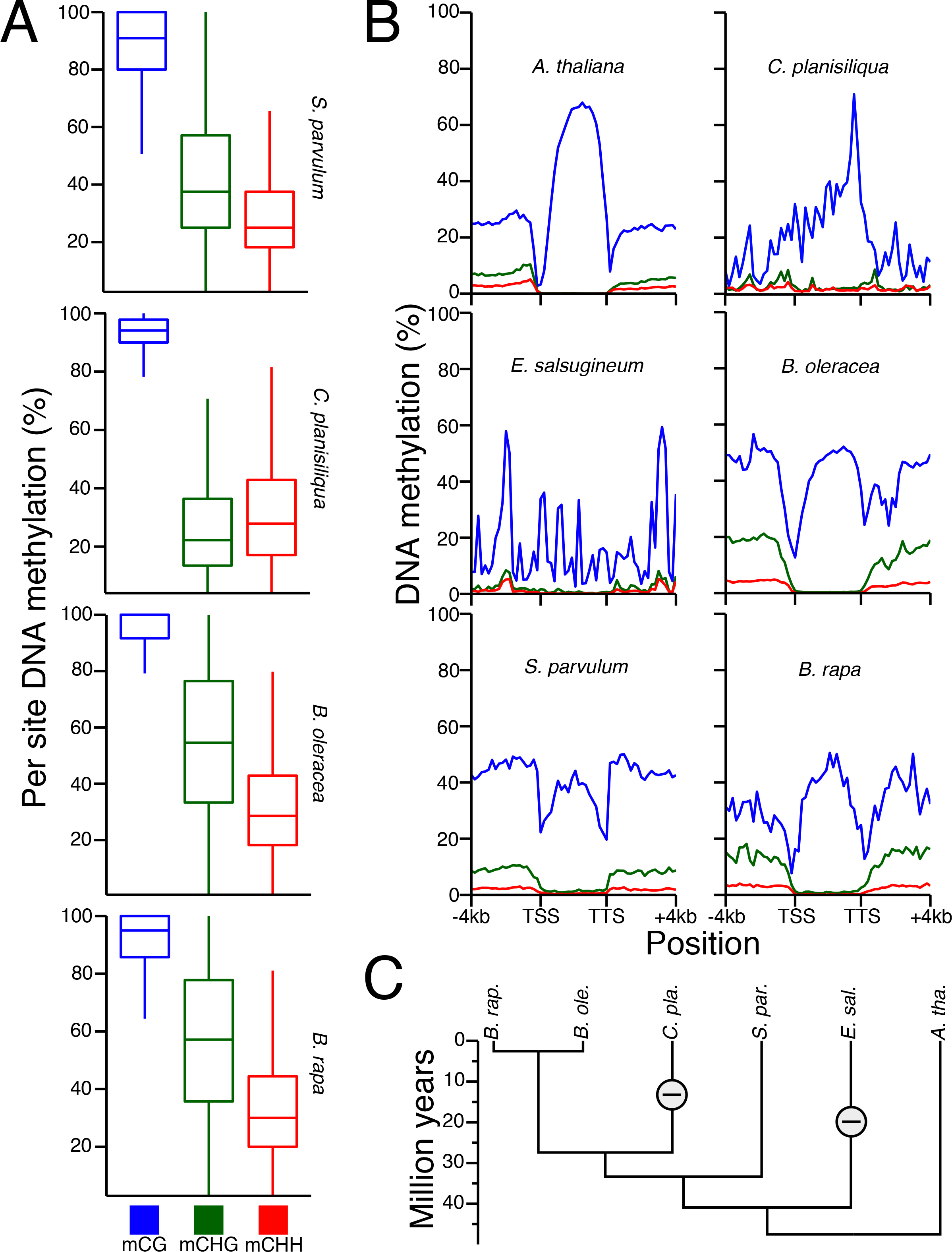
**CMT3 is required for CHG DNA methylation and establishment of gbM in angiosperms**. **(A)** Similar to *E. salsugineum*, loss of CMT3 in *C. planisiliqua* leads to genome-wide reductions of CHG DNA methylation at a per site level. Closely related species that possess CMT3 maintain higher per site CHG DNA methylation compared to *C. planisiliqua*. **(B)** The loss of CMT3 also causes the loss of gene body methylation in *C. planisiliqua*. **(C)** The loss of CMT3 (-) in *E. salsugineum* and *C. planisiliqua* represents two independent events separated by at least 13.5 MY; loss of CMT3 in *C. planisiliqua* and *E. salsugineum* occurred within the last 27.41 MY and 40.92, respectively (20). However, loss in *C. planisiliqua* could be more recent; ≤12.27 MY (20).

The methylome of *E. salsugineum*, with lower mCHG levels in repeats and a complete loss of gbM was unique compared to 86 *A. thaliana* mutants for which methylome data were available (18). The absence of CMT3 from *E. salsugineum* is consistent with two characteristics of mCHG in its genome. First, the methylation level at individual CHG sites was significantly lower than any other species and was similar to the *A. thaliana cmt3* mutant (**Fig. 1D**). Second, as CHG is symmetrical, with a mirrored cytosine on the opposing strand, CMT3 activity results in high methylation of cytosines on both strands. In *E. salsugineum* the percentage of paired CHG sites that were highly methylated is significantly lower than wild-type *A. thaliana* and again similar to the *cmt3* mutant, suggesting that the mCHG in *E. salsugineum* is likely a result of RdDM activity (**Fig. 1E** and **F**). Taken together, these results indicated that *E. salsugineum* does not have CMT3 activity.

The loss of *CMT3* and gbM from *E. salsugineum* is consistent with the hypothesis that CMT3 is required for the establishment of gbM. To solidify this connection we searched for additional angiosperms that do not possess CMT3. Curiously, we identified another Brassicaceae, *Conringia planisiliqua*, which is also missing CMT3. Methylome analysis of *C. planisiliqua*, and other closely related Brassicaceae *(Brassica rapa, Brassica oleracea*, and *Schrenkeilla parvula*) which all possess a CMT3 confirmed the presence of CHG methylation typical of CMT3 activity (Fig. 2A). However, the CHG methylation present in *C. planisiliqua* was similar to that observed in *E. salsugineum* and *cmt3* mutants (**Fig. 1D**), indicating that the CHG methylation detected is likely a result of RdDM and not maintenance by CMT3. Loci containing gbM were identified using the same methods defined previously (see methods) and CG, CHG and CHH methylation metagene plots of these defined loci were generated for each of these Brassicaceae (**Fig. 2B**). All of the species that possess a functional CMT3 also possess gbM, whereas the two species that do not possess CMT3 (*E. salsugineum* and *C. planisiliqua*) do not possess patterns consistent of gbM loci (**Fig. 2B**). In addition, metagene plots of CG methylation across all genes reveal a complete absence of gbM in *C. planisiliqua* (**Fig. S2**), which is similar to observations in *E. salsugineum* (**Fig 1A**). Therefore, given the evolutionary relationship between these Brassicaceae species (**Fig. 2C**), the most parsimonious explanation is two independent losses of CMT3. Additionally, synteny between *A. thaliana*, *C. planisiliqua*, and *E. salsugineum* supports the hypothesis of unique events that led to the deletion of CMT3 in *C. planisiliqua* and *E. salsugineum* (**Fig. S3**). Taken together, these results strongly suggest that CMT3 is involved in the establishment of gbM in angiosperms.

In addition to the identification of CMT3 as the enzyme correlated with the establishment of gbM, the methylome data generated here also provided insights into why only a subset of the genes in each plant genome contained gbM. Previous studies have shown that gbM genes tend to be expressed at moderate to high levels in *A. thaliana* (4, 5, 9). Similar results were found in all other plant species that contained gbM (19). These results indicated that the CMT3/KYP pathway might preferentially target moderately to highly transcribed genes. The mechanistic basis for this preference is not yet clear. It is possible that H3K9me2-containing H3 may be occasionally mis-incorporated into active genes during transcription-coupled nucleosome turnover. Alternatively, nucleosome movement during transcription may lead to the spreading of mCHG/H3K9me2 in flanking regions into active genes, as was described at the *A. thaliana BONSAI* gene (19).

A significant fraction of genes expressed at moderate to high levels did not contain gbM, indicating that active transcription in itself may not be sufficient to trigger gbM. A comparison of the DNA sequence content in gbM and UM (unmethylated) genes expressed at comparable expression levels (**Fig. S4**) revealed a difference in the frequency of CAG/CTG sites per kilobase pair of gene length; gbM loci had higher densities of these sites at a frequency of 60.2/kb compared to 45.4/kb for UM genes. These results indicate that gbM loci are potentially predisposed to accumulation of gbM because of their base composition in combination with transcriptional activity.

To obtain further evidence regarding the role of active transcription and sequence composition in the establishment of gbM, we took advantage of the availability of 8^th^ generation *met1* epigenetic recombinant inbred lines (epiRILs) (21), and determined the characteristics of genes where gbM was reestablished. The epiRILs contain mosaic methylomes, including genomic regions with normal methylation from the wild-type parent, and gbM-free regions from the *met1* parent. As there was very limited genetic variation between the wild-type and *met1* parents, we determined the parental origin of each genomic region according to gbM patterns, similar to what was previously performed for the *ddm1* epiRIL population (**Fig. 3A** and **Fig. S5**) (22, 23). Although mCG was reestablished in repetitive sequences in *met1-derived* regions, the vast majority of genes remained mCG-free, indicating that in most cases gbM has not returned (**Fig. 3B** and **C**). A closer inspection identified some rare loci where gbM was partially restored. A total of 50, 10, and 29 genes were identified that accumulated >5% mCG methylation *met1* epiRIL–1, −12 and −28 lines, respectively (**Fig. 3D**). The loci where mCG returned rarely overlapped in the three epiRILs, indicating that the establishment of gbM is a slow and stochastic process. Consistent with the results described earlier, the genes with partially restored gbM tend to be moderately expressed and possess higher frequencies of CAG/CTG sites, indicating that genes with active transcription and higher densities of CAG/CTG sites might be more susceptible to the establishment of gbM.

**Fig. 3.**
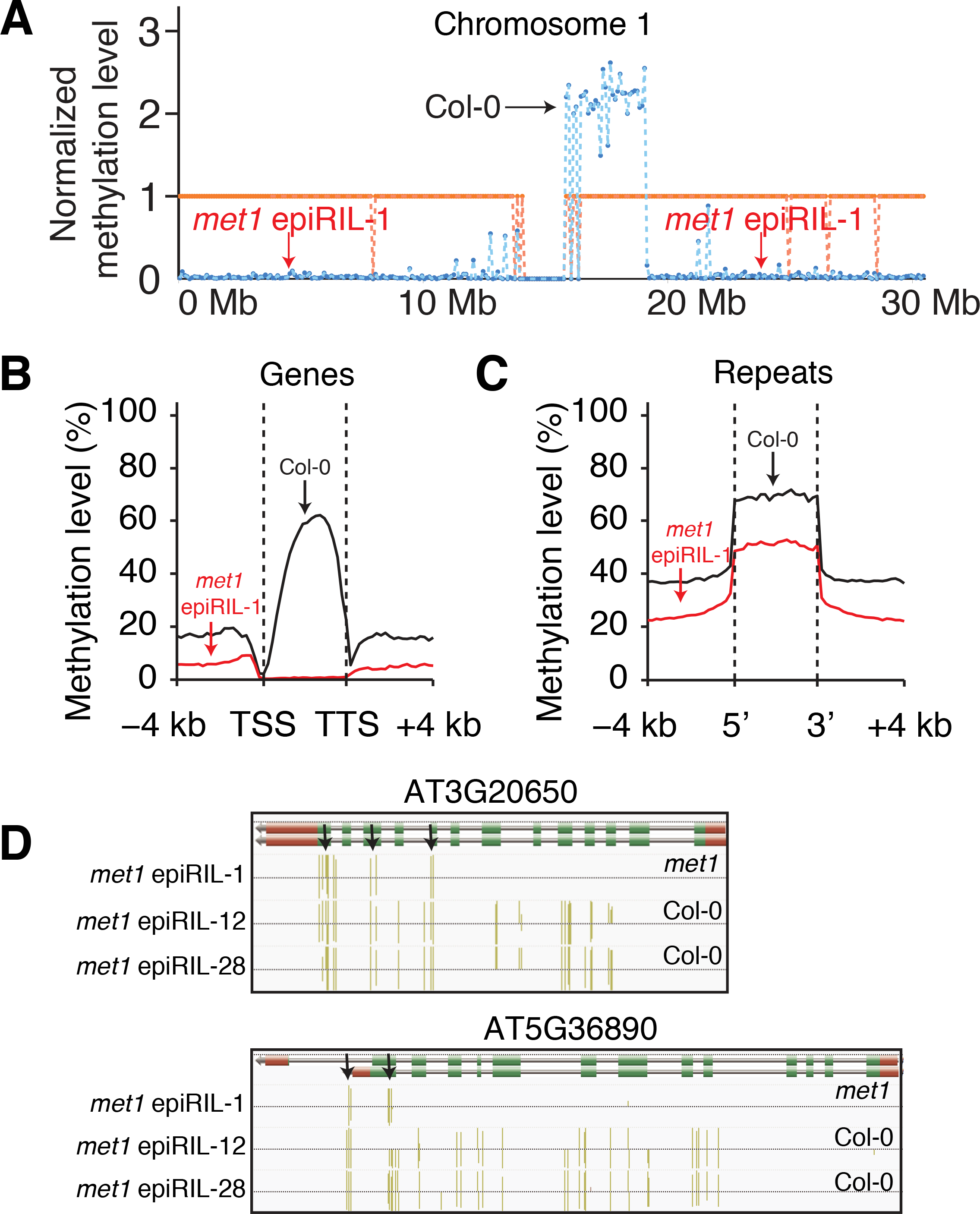
***De novo* gbM accumulates incrementally over generational time**. **(A)** A genetic map of the *met1* epiRIL line 1 using only the methylation status of gbM loci from wild-type Col–0 as markers. The orange midpoint line indicates the heterozygous methylation levels from Col-0 and the blue line indicates observed results from the *met1* epiRIL that was analyzed. Metaplots of CG methylation in **(B)** genes and **(C)** transposons including 4kb up-and down-stream of the TSS (transcriptional start site) and TTS (transcriptional termination site) from the Col–0 or the *met1* derived regions of the epiRIL. **(D)** Examples of loci in *met1* epiRIL–1 where gbM has partially returned in loci that are located in *met1* derived regions of the genome. Black arrows indicate where mCG returned.

gbM has been proposed to function in several steps in gene expression, such as suppressing antisense transcription, impeding with transcriptional elongation and thus negatively regulating gene expression, or affecting post-transcriptional RNA processing such as splicing (5, 9, 24, 25). However, comparisons between RNA-seq data from *met1* epiRILs and wild-type Col–0 revealed no evidence in support of these possibilities (**Table S2–4**).

It is possible the function of gbM has been masked by redundant contributions from other chromatin modification pathways. For example, H3K36me3 has been shown to function in suppressing cryptic transcriptional initiation and regulating splicing (26). We therefore created the triple mutant *met1 sdg7 sdg8*, in which both gbM and H3K36me3 were eliminated (27). RNA-seq experiments from *met1 sdg7 sdg8* and wild-type Col–0 again failed to identify significant differences in mRNA expression, antisense transcription, or splicing variants in a comparison between gbM loci and UM loci (**Table S5–7**).

The effect of gbM on gene expression might only become detectable over an evolutionary time scale. The identification of plant species with no gbM provided a unique opportunity to test this possibility. We determined the gene set in the *E. salsugineum* genome that likely contained gbM before the loss of CMT3 by projecting orthologous *A. thaliana* gbM genes onto *E. salsugineum*, as they may have at one point possessed gbM in a common ancestor. RNA-seq analysis showed that *E. salsugineum* genes predicted to have contained gbM exhibited similar transcription levels to their *A. thaliana* orthologs (**Fig. 4A**). These results suggest that gbM may have a limited role in transcriptional regulation.

**Fig. 4.**
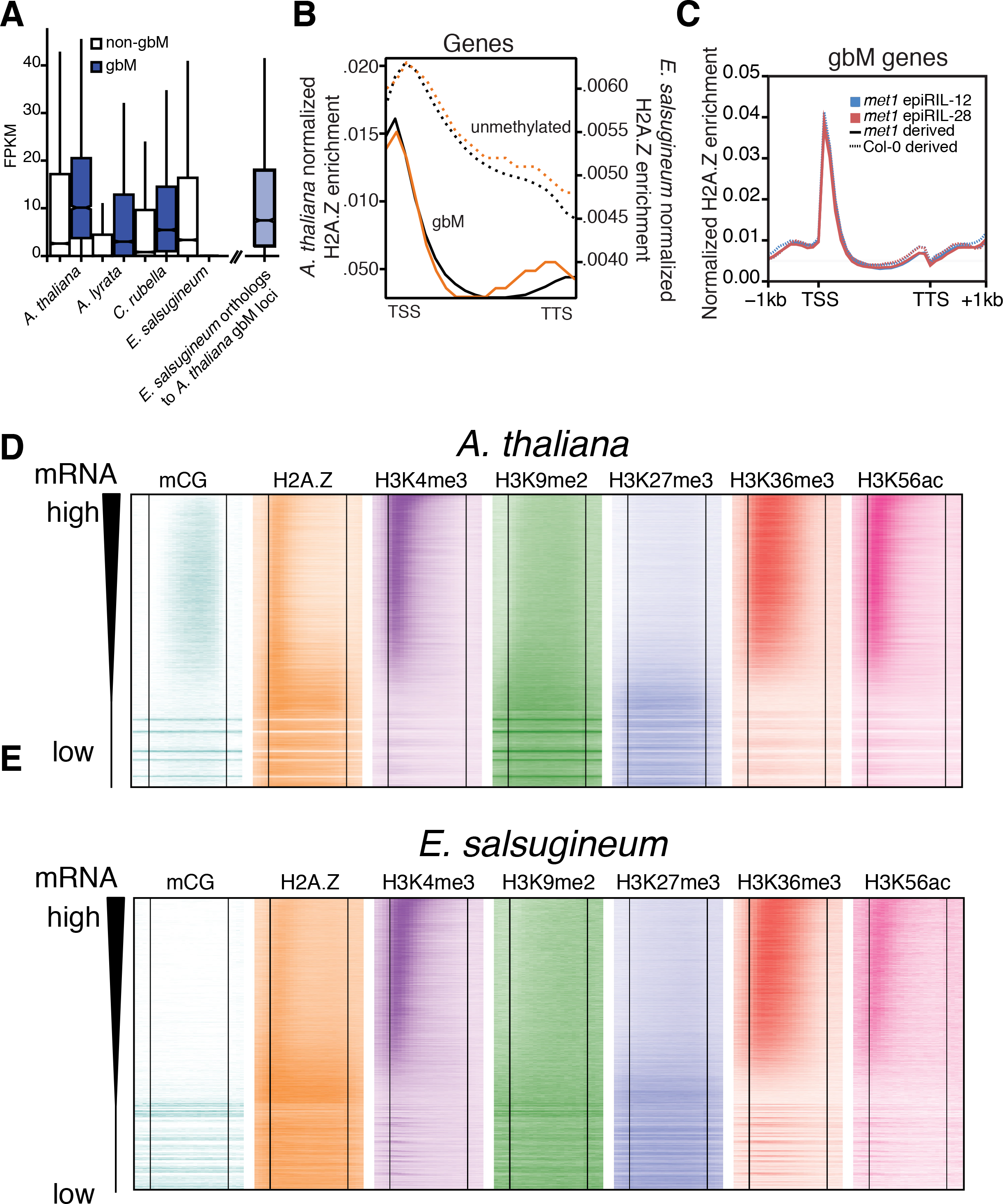
**Gene expression and histone modifications are not affected by loss of gbM in *E. salsugineum***. **(A)** Comparison of gene expression levels between gbM and non-gbM within each species listed. Orthologs of gbM loci from *A. thaliana* were used to identify loci from *E. salsugineum* and FPKM values were plotted. **(B)** Metagene plots of H2A.Z enrichment in gbM versus unmethylated genes in *A. thaliana* (black line) and *E. salsugineum* (orange line). The y-axis on the left is associated with *A. thaliana* and the one on the right is associated with *E. salsugineum*. **(C)** Metagene plots of H2A.Z enrichment in two *met1* epiRILs of gbM loci derived from either the Col–0 (dotted lines) or the *met1* (solid lines) parent. Heatmap representation of histone modification distributions and patterns in gene bodies of **(D)** *A. thaliana* and **(E)** *E. salsugineum.* The genes in each heatmap are ranked from highest to lowest expression levels. The vertical lines indicate the position of the transcriptional start and stop sites. 1 kb upstream of the TSS and downstream of the TTS is included in the heatmaps.

In addition to gene expression, gbM has also been proposed to function to prevent the histone variant H2A.Z from encroaching into gene bodies (9, 24). To test this hypothesis, chromatin immunoprecipitation-sequencing (ChIP-seq) was performed using an H2A.Z antibody on chromatin isolated from *E. salsugineum* and *A. thaliana* leaves. The distribution pattern of H2A.Z in *A. thaliana* was found to be highly consistent with previously published results: in gbM genes, H2A.Z was enriched at the 5’ ends and absent from regions within genes that contained gbM; in non-gbM genes, H2A.Z could be found distributed throughout the gene bodies (**Fig. 4B**). Although gbM is presumably absent from *E. salsugineum* for a considerable amount of time, the distribution pattern of H2A.Z remained comparable to that in *A. thaliana* (**Fig. 4B**). Measuring enrichment of H2A.Z in two independent *met1* epiRILs revealed a similar result. The distribution and amplitude of H2A.Z was similar at a previously defined set of gbM loci regardless of whether the gene was inherited from the Col–0 or the *met1* parental genome (**Fig. 4C**). The mechanistic basis for the correlation between H2A.Z and transcription levels is not clear. Regardless, these results showed that the loss gbM over an evolutionary time scale has no effect on the distribution of H2A.Z.

The loss of gbM in *E. salsugineum* might be compensated by a redistribution of other histone modifications across gene bodies. Therefore, additional ChIP-seq experiments using antibodies against H3K4me3, H3K9me2, H3K27me3, H3K36me3, and H3K56ac were performed to test if the absence of gbM in *E. salsugineum* affected the distribution of these histone modifications. However, no differences in distributions were observed when compared against *A. thaliana* (**Fig. 4D and E**).

We propose that gbM might represent a byproduct of errant properties associated with enzymes that can establish DNA methylation, like CMT3, and enzymes that can maintain it, such as MET1. Loss of IBM1, a histone demethylase, results in immediate accumulation of H3K9me2 and CHG methylation in gene bodies (16, 17). In fact, DNA methylome profiling of an *ibm1-6* allele not only confirmed these results, but also uncovered an increase of both CG and CHH methylation in gbM loci (**Fig. S6**). This indicates that in the absence of active removal of H3K9me2 from gene bodies, methylation in all cytosine sequence contexts accumulates. Failure to properly remove H3K9me2 accumulation from gene bodies of wild-type plants leads to recruitment of CMT3, which in turn methylates cytosines primarily in the CHG context, but also enables methylation of CG and CHH sites (**Fig. S6**). Once methylation is present in the gene bodies, it spreads throughout the gene body, as methylated DNA serves as a substrate for the SRA-domain containing proteins KYP/SUVH4/SUVH5, which binds methylated cytosines and leads to continual methylation of H3K9 (28). The lack of gbM at the TSS might be due to an inability of H2A.Z and H3K9me2 to cooccur in nucleosomes, which suggests the primary role of H2A.Z is to prevent spreading of H3K9me2 into the TSS. This spreading mechanism is also consistent with the loci in the *met1* epiRILs, where gbM partially returned seemingly in a directional manner (**Fig. 3D**). Therefore, over evolutionary time scales, gbM accumulates and is maintained and tolerated by the genome, as thus far it appears to have no apparent functional role and no deleterious consequences.

It cannot be proven that gbM has no functional role in angiosperm genomes, as it is possible that it has a yet undiscovered function or that it serves to redundantly perform functions with other transcriptional processes. However, the absence of gbM in *E. salsugineum* and *C. planisiliqua* and their perseverance as a species is clear evidence that this feature of the DNA methylome is not required for viability. DNA methylation of gene bodies is also found in mammalian genomes, although its distribution throughout transcribed regions and its mechanism for establishment is distinct from angiosperms. In mammals, the methyltransferase Dnmt3B binds H3K36me3 through its PWWP domain and catalyzes *de novo* methylation in gene bodies, which is then maintained by the maintenance methyltransferase Dnmt1 (29). Intriguingly, loss of methylation in gene bodies in a mouse methylation mutant did not result in global reprogramming of the transcriptome, as the correlation of mRNA levels between mutant and wild type was 0.9611 (30). Only a handful of differentially expressed loci were found. Therefore, although there is abundant DNA methylation of gene bodies in mammals and certain plant species, the evidence available thus far does not provide strong support for a functional role in gene regulation. Future studies of this enigmatic feature of genomes will be required to understand whether gene body methylation has a function or is simply a byproduct of active DNA methylation targeting systems.

## Author Information

Analyzed datasets can be obtained from http://schmitzlab.genetics.uga.edu/plantmethylomes. Sequence data for MethylC-seq, RNA-seq and ChIP-seq experiments are available at NCBI Gene Expression Omnibus under the accession number GSE75071. Correspondence and requests for materials should be addressed to R.J.S.

## Methods

**Plant material.** Leaf tissue was flash frozen in liquid nitrogen for all experiments including ChIP-seq, RNA-seq and MethylC-seq. DNA was isolated using a Qiagen Plant DNeasy kit (Qiagen, Valencia, CA) following the manufacturer's recommendations. RNA was isolated using TRIzol (Thermo Scientific, Waltham, MA) following the manufacturer's instructions. Specimens of *Conringia planisiliqua* (B–2011–0093, HEID921022) were cultivated as described below. We collected the 11^th^ leaf from two plants resulting in two biological replicates. The DNA was isolated from the young leaf tissue flash frozen in liquid nitrogen using a Plant DNeasy Mini Kit from Qiagen following the manufacturer's protocol.

**Sequencing, assembly and annotation of *C. planisiliqua* gnome**. Specimens of *Conringia planisiliqua* (B–2011–0093, HEID921022, kindly provided by Marcus Koch, University of Heidelberg/Germany) were grown in the greenhouse (temperature day 20°C/night 18°C; cycles of 16h light and 8h darkness, if required day length was extended to 16h by additional light sources (Osram HQI-BT 400W/D (white), NAV400W (orange/red)) or alternatively plants were shaded from light to keep day length constant. Plants were cultivated on soil (Balster, Type Mini Tray with 1kg/m^3^ added fertilizer and 1kg/m^3^ Osmocote (Scouts)), watered daily and fertilized weekly with Wuxal Super 8–8–6.

A whole genome shotgun library with a 300 bp insert size was generated following the manufacturer's protocol and sequenced paired-end on an Illumina HiSeq2500 with 101 bp read lengths. We obtained 28,942,646 read-pairs. A genome size estimation based on kmers indicated a genome size of 260 Mb, which is slightly higher than what has been estimated based on flow cytometry (180 Mb). We generated a whole genome assembly using Platanus (v1.2, (31)). The final assembly consisted of 12,970 scaffolds larger than 500 bp with L50 of 37 kb and a N50 of 729 summing up to a total length of 121 Mb. For homology-based annotation we aligned the protein sequences of *A. thaliana, A. lyrata, S. parvula, E. salsugineum, B. rapa, A. alpina* and *A. arabicum* against the scaffolds using scipio (1.4, (32)). The blat alignment files were filtered using a perl script (“filterPSL.pl”) provided with Augustus v3.0 (33) using minCover=80 and minId=80 as parameters. The filtered alignment file was than converted into gff format using another perl script (“blat2hints.pl”) provided with Augustus and was used as input for Augustus *de novo* gene prediction (v3.0.1) using the adapted training parameters for *Arabidopsis* (--species=arabidopsis). Augustus predicted 29,373 genes. In order to check for completeness of our gene set, we blasted the protein sequences against the set of core eukaryotic genes of *A. thaliana* (34) (http://korflab.ucdavis.edu/datasets/cegma/#SCT2) and found for 455 out of 458 a blast hit with an e-value <1e-50 and an identity >70%. Reciprocal best BLAST was performed between *C. planisiliqua* protein coding genes and *A. thaliana* transposable elements (TEs) (TAIR10). Using an e-value cutoff of ≤1e–06 we identified 795 TEs annotated as protein coding genes; these TEs were removed prior to any analyses relating to gbM.

**MethylC-seq library construction**. All libraries were prepared as described by Urich et al (35) with the exception of *Conringia planisiliqua*. Genomic DNA (gDNA) was quantified using the Qubit BR assay (ThermoFisher Scientific, U.S.A.) and the quality assessed by standard agarose gel electrophoresis. Bisulfite libraries were constructed using the NEXTflex Bisulfite-Seq Kit (U.S.A) with 500ng input gDNA being fragmented by COVARIS S2 and then increased in concentration with AMPure Beads (0.8 volume, Beckman Coulter, U.S.A) and eluted in water. Sheared Lambda DNA was spiked as to record the bisulfite conversion rate. For ligation, the adapter concentration was diluted 1:1. Library fragments were then two times purified with 1 volume AMPure Beads and finally eluted in 10mM Tris pH8. Bisulfite conversion of library fragments was performed as outlined in the EZ DNA Methylation-Gold Kit (Zymo Research Corporation, U.S.A.). Converted DNA strands were amplified with 15 PCR cycles, purified with AMPure beads followed by quality assessment with capillary electrophoresis (D 1000 Bioanalyser Assay, Agilent, U.S.A) and quantified by fluorometry (Qubit HS assay; ThermoFisher Scientific, U.S.A.).

**ChIP-seq library construction**. ChIP experiments were performed as described in (36). Immunoprecipitated DNA was end repaired using the End-It DNA Repair Kit (Epicentre, Madison, WI) according to the manufacturer's instructions. DNA was purified using Sera-Mag (Thermo Scientific, Waltham, MA) at a 1:1 DNA to beads ratio. The reaction was then incubated for 10 minutes at room temperature, placed on a magnet to immobilize the beads, and the supernatant was removed. The samples were washed two times with 500μl of 80% ethanol, air dried at 37°C and then resuspended in 50μl of 10 mM Tris-Cl pH8.0. Finally, the samples were incubated at room temperature for 10 minutes, placed on the magnet, and the supernatant was transferred to a new tube, which contained reagents for “A-tailing.” A-tailing reactions were performed at 37°C according to the manufacturer's instructions (New England Biolabs, Ipswich, MA). The samples were cleaned using Sera-Mag beads as previously described. Next, adapter ligation was performed using Illumina Truseq Universal adapters and T4 DNA ligase (New England Biolabs, Ipswich, MA) overnight at 16°C. A double clean-up using the Sera-Mag beads was performed to remove any adapter-adapter dimers and the elution was used for 15 rounds of PCR. Lastly, samples were cleaned up one final time using the procedures described above.

**RNA-seq library construction**. RNA-seq libraries were constructed using Illumina TruSeq Stranded RNA LT Kit (Illumina, San Diego, CA) following the manufacturer's instructions with limited modifications. The starting quantity of total RNA was adjusted to 1.3μg, and all volumes were reduced to a third of the described quantity.

**Sequencing**. MethylC-Seq libraries for the species survey were sequenced using the Illumina HiSeq 2500 (Illumina, San Diego, CA). Sequencing of libraries was performed up to 101 cycles. Wild-type, *met1* epiRILs and *Ibm1–6* methylomes and transcriptomes were sequenced to 150 bp on an Illumina NextSeq500 (Illumina, San Deigo, CA) with the exception of wild-type and *sdg7/sdg8/met1* transcriptomes, which were sequenced using the Illumina HiSeq2000 (Illumina, San Diego, CA).

**MethylC-seq analysis**. Genomes and annotations were downloaded from Phytozome 10.3 (http://phytozome.jgi.doe.gov/pz/portal.html) (37). Transposons annotations for *A. thaliana* were downloaded from TAIR (https://www.arabidopsis.org/). MethylC-seq reads were processed, aligned, and sites called using published methods (38). The genome is converted into a forward strand reference (all Cs to Ts) and a reverse strand reference (all Gs to As). Cutadapt 1.1.0 (39) is used to trim adaptor sequences and then bowtie 1.1.0 (40) aligns reads to the two converted reference genomes. Only uniquely aligned reads are retained and the non-conversion rate calculated from unmethylated reads aligned to the chloroplast genome or spiked in unmethylated lambda DNA. Summary statistics for each methylome sequenced are present in Table S1. A binomial test is then applied to each cytosine and corrected for multiple testing using Benjamini-Hochberg False Discovery Rate. A minimum of three reads is required at each site to call it as methylated. For the *cmt3–11t* and *met1–3* methylation data, raw reads were downloaded from a previously published study (18).

**Metagene plots**. The gene body was divided into 20 windows. Additionally, regions 4000 bps upstream and downstream were each divided into 20, 200 bp windows. Weighted methylation levels were calculated for each window (41). For gene bodies, only cytosine sites within coding sequences were included. For repeat bodies, all cytosine sites were included. The mean weighted methylation for each window was then calculated for all genes and plotted in R.

**Analysis of mCHG characteristic of CMT3 activity**. To create the plots for **Figure 1F** both strands of each CHG sequence were required to have a minimum coverage of at least three reads and at least one of the CHG sites was identified as methylated as described previously in the MethylC-seq analysis section. Methylation levels were calculated for each strand of the symmetric CHG site and values used to create a density plot using R. To perform the analysis presented in **Figure 1E**, both strands of each CHG dinucleotide were required to have a minimum coverage of at least three reads and at least one of the CHG sites was identified as methylated.

**Identification of CG gbM genes**. Genes were identified as gbM (**Dataset S1**) using a modified version of the binomial test used by Takuno and Gaut (42). This approach tests for enrichment of CG, CHG, and CHH against a background level calculated from the entire set of genes. To do this, the total number of cytosines in each context (CG, CHG, CHH) with mapped reads and the total number of methylated cytosines called were calculated for the coding regions of each gene. Because species differ in genome size, TE content, and other factors; this analysis was restricted to coding sequences and a single universal background level of methylation was calculated by combining data from all species and determining the percentage of methylated cytosines in each context for the coding regions. A one-tailed binomial test was then applied to each gene for each context testing against the background methylation level. Tested across tens of thousands of genes, there will be some false positives at the extremes. To control for this type I error, q-values were calculated from p-values by adjusting for multiple testing using the Benjamini-Hochberg False Discovery Rate. GbM genes were defined as having reads aligning to at least 20 CG sites and a CG methylation q-value <0.05 and CHG and CHH methylation q-values >0.05. Thus, this test identifies gene-coding sequences that are enriched for mCG, and depleted in mCHG and mCHH.

**RNA-seq data analysis**. Raw FASTQ reads were trimmed for adapters, preprocessed to remove low quality reads using Trimmomatic v0.32 (43). Reads were aligned using Tophat v2.0.13 (44) supplied with a reference GFF file and the following arguments: −I 50000––b2-very-sensitive––b2-D 50. Transcripts were then quantified using Cufflinks v2.2.1 supplied with a reference GFF file (45). For the *A. thaliana* samples, rRNA contaminants were removed using an *A. thaliana* rRNA database (46) using BLAT v35 (47). Differentially expressed genes were determined by Cufflinks requiring statistically significant changes and also by requiring a 2–fold (log2) change in gene expression.

**Intron retention analysis**. The *A. thaliana* TAIR10 GFF file was downloaded and intron entries were created. All gene annotation entries were removed except ‘gene’, ‘mRNA’, and ‘intron’. Then introns were renamed to exons prior to using TopHat and Cufflinks. Gene expression values and detection of differentially expressed genes were determined using the same process and criteria as we described above for the detection of differentially expressed mRNAs.

**Antisense transcription analysis**. Using.bam files generated from RNA-seq alignments as described previously, reads that mapped to convergently transcribed genes were removed from all subsequent analysis. The TAIR10 gff file was modified to generate an “antisense transcription annotation gff file” by reversing strand orientation of all annotated features. Using this file coupled with the filtered.bam files we determined the prevalence of antisense transcription using the same process and criteria as we described above for the detection of differentially expressed mRNAs.

**Synteny analysis between *A. thaliana* and *E. salsugineum***. Whole-genome synteny between *A. thaliana* and *E. salsugineum* was determined using CoGe's SynFind program (48). *A. thaliana* chromosome 1, which harbors CMT3, and *E. salsugineum* scaffold 9 were identified as being syntenic. Finer synteny analysis was performed on *A. thaliana* chromosome 1 and 150 kb upstream and downstream of CMT3, and *E. salsugineum* scaffold 9 using CoGe's GEvo program (48).

**Identifying orthologs and estimating evolutionary rates**. Reciprocal best BLAST with an e-value cutoff of ≤1e-08 was used to identify orthologs. Individual protein pairs were aligned using Multiple Sequence Comparison by Log-Expectation (MUSCLE) (49), and back-translated into codon alignments using the CDS sequence. Insertion-deletion (indel) sites were removed from both sequences, and the remaining sequence fragments were concatenated into a contiguous sequence. A ≥ 30 bp and ≥ 300 bp cutoff for retained fragment length after indel removal, and concatenated sequence length was implemented, respectively. Subsequently, substitution rates were calculated for sequence pairs between *A. thaliana* and *E. salsugineum* using the yn00 method implemented in Phylogenetic Analysis by Maximum Likelihood (PAML) (50).

**Analysis of a *metl* epiRIL methylome and generation of epigenetic maps**. Each chromosome was split every 100 kb and weighted methylation levels were computed from gbM loci in each bin for each *met1* epiRIL and Col–0. Only cytosines in coding regions were used to compute methylation levels. For each bin, the midpoint of methylation level in Col–0 was defined as the heterozygous methylation level. Methylation levels from each *met1* epiRIL sample were divided by heterozygous methylation levels to normalize each bin. Horizontal line (Y=1) was defined as the heterozygous line and bins without any gbM loci were not assigned any values. Normalized *met1* epiRIL methylation levels above the heterozygous line were regarded as being derived from the Col–0 parent, whereas data points below the line were regarded as being derived from the *met1* parent. Bins located near the crossover events were excluded from subsequent analyses.

**ChIP-seq data analysis**. Raw ChIP reads were trimmed for adapters and low-quality bases using Trimmomatic version 0.32. Reads were trimmed for TruSeq version 3 single-end adapters with maximum of two seed mismatches, palindrome clip threshold of 30 and simple clip threshold of 10. Additionally, leading and trailing bases with quality less than 10 were removed; reads shorter than 50 bp were discarded. Trimmed reads were mapped to the TAIR10 genome using bowtie2 version 2.2.3 with default options. Mapped reads were sorted using samtools version 1.2 then clonal duplicates were removed using samtools version 0.1.9. Remaining reads were converted to BED format with bedtools version 2.21.1.

**Generation of H2A.Z heatmaps and metagene plots**. For heatmaps and metagene plots, intergenic regions, defined as the region between genes excluding 100 bp upstream and downstream of genes, were determined for both *A. thaliana* and *E. salsugineum* using the TAIR10 and Phytozome 10 *E. salsugineum* 173 v1 annotations, respectively. Intergenic regions were broken into 2000 bp segments and 25000 segments with at least 40 CpG sites were randomly chosen. Segments were ranked by weighted methylation in all contexts. The 2000 segments with highest methylation and 2000 segments with lowest methylation were defined as intergenic methylated and intergenic unmethylated, respectively. Orthologs between *A. thaliana* and *E. salsugineum* were split into two groups, gbM and UM as defined by the methylation in *A. thaliana*. Coordinates for the genes were taken from the TAIR10 annotation of *A. thaliana* and Phytozome 10 annotation of *E. salsugineum 173 v1*. All intergenic segments and orthologs were broken into 20 equally sized bins. Aligned ChIP reads starting the each bin were summed then normalized by bin length and non-clonal library size.

For heatmaps, the 95^th^ percentile value of all bins was computed and any bin with value above this threshold was set equal to the threshold. Finally, the average bin value for bins of intergenic methylated regions was computed. This value was subtracted from all bins in the heatmap and any bin value less than zero was set equal to zero. Orthologs were ordered based on mRNA level in *A. thaliana*. For metagene plots, bin values were summed for each ortholog type, gbM and UM then normalized for the number of genes in each group.

## Acknowledgements

We thank Zachary Lewis, Nathan Springer and Dave Hall for comments and discussions as well as Karen Schumaker (*E. salsugineum*), Marcus Koch (*C. planisiliqua*) and Jerzy Paszkowksi *(met1* epiRILs) for seeds and the Georgia Genomics Facility and the Georgia Advanced Computing Resource Center for technical support. This work was supported by the National Institutes of Health (R00GM100000), by The Pew Charitable Trusts and by the Office of the Vice President of Research at UGA to R.J.S. C.E.N was supported by a National Science Foundation (NSF) postdoctoral fellowship (IOS – 1402183). Research in X.Z. laboratory was supported by NSF (MCB – 0960425). B. T.H. was supported by a Scholars of Excellence Graduate Fellowship from UGA. H.S. was supported by an HHMI Fellow of the Damon Runyon Cancer Research Foundation (DRG–2194–14). S.E.J. is an Investigator of the Howard Hughes Medical Institute.

